# The malaria-protective human glycophorin structural variant DUP4 shows somatic mosaicism and association with hemoglobin levels

**DOI:** 10.1101/360453

**Authors:** Walid Algady, Sandra Louzada, Danielle Carpenter, Paulina Brajer, Anna Färnert, Ingegerd Rooth, Fengtang Yang, Marie-Anne Shaw, Edward J Hollox

## Abstract

Glycophorin A and glycophorin B are red blood cell surface proteins that are both receptors for the parasite *Plasmodium falciparum*, which is the principal cause of malaria in sub-Saharan Africa. DUP4 is a complex structural genomic variant that carries extra copies of a glycophorin A - glycophorin B fusion gene, and has a dramatic effect on malaria risk by reducing the risk of severe malaria by up to 40%. Using fiber-FISH and Illumina sequencing, we validate the structural arrangement of the glycophorin locus in the DUP4 variant, and reveal somatic variation in copy number of the glycophorin A-glycophorin B fusion gene. By developing a simple, specific, PCR-based assay for DUP4 we show the DUP4 variant reaches a frequency of 13% in a village in south-eastern Tanzania. We genotype a substantial proportion of that village and demonstrate an association of DUP4 genotype with hemoglobin levels, a phenotype related to malaria, using a family-based association test. Taken together, we show that DUP4 is a complex structural variant that may be susceptible to somatic variation, and show that it is associated with a malarial-related phenotype in a non-hospitalized population.

**Significance statement:** Previous work has identified a human complex genomic structural variant called DUP4, which includes two novel glycophorin A-glycophorin B fusion genes, is associated with a profound protection against severe malaria. In this study, we present data showing the molecular basis of this complex variant. We also show evidence of somatic variation in the copy number of the fusion genes. We develop a simple robust assay for this variant and demonstrate that DUP4 is at an appreciable population frequency in Tanzania and that it is associated with higher hemoglobin levels in a malaria-endemic village. We suggest that DUP4 is therefore protective against malarial anemia.

## Introduction

Structural variation (SV) of genomes, including inversions, deletions, duplications and more complex rearrangements is seen at polymorphic frequencies across all species. Like single nucleotide variation, much SV is likely to be evolving neutrally but in some cases there is evidence for balancing or adaptive evolution (1–5). SV has also been shown to generate novel genes with functional consequences, for example generation of human-specific *SRGAP2* genes that increase the density of dendritic spines in the brain (6, 7). Regions that show extensive SV are thought to have a high mutation rate, due to recurrent non-allelic homologous recombination (8–11). In addition to SV in the germline, large somatic SVs have been observed in human brain (12), skin fibroblasts (13), and in blood of identical twins (14).

Although SV is a source of variation between individuals (15), and between cells within an individual (16), its contribution to disease resistance remain unclear, and the best-characterized examples of structural variants associated with a common disease involve identifying a germline structural variant in linkage disequilibrium with a sentinel single-nucleotide variant (SNV) that has been previously highlighted in a large genome-wide association study (17).

The human genome assembly carries three glycophorin genes, *GYPE, GYPB* and *GYPA*, tandemly arranged on three ~120 kb repeats sharing ~97% identity. Glycophorin A (encoded by *GYPA*) and Glycophorin B (encoded by *GYPB*) are readily detectable on the surface of erythrocytes, and carry the MNS blood group system (18). Mature glycophorin E (encoded by *GYPE*) is predicted to be 59 amino acids long but has not been unambiguously detected on the erythrocyte surface. This genomic region carrying this genes is known to undergo extensive copy number variation and gene conversion, and rearrangements that shuffle the coding regions of the three genes can generate rare blood group antigens in the Miltenberger series (19). This genomic region has also been highlighted as a region of balancing selection (20–23) and positive selection (24), though the effect of extensive gene conversion between the 120kb repeats on statistical measures of positive selection is not clear.

Recent work studying the host genetic contribution to severe malaria identified a SNV allele at a nearby non-repeated region in linkage disequilibrium with a complex structural variant at the glycophorin locus protective against severe malaria (5, 20). This structural variant, called DUP4, is restricted to East African populations, and is responsible for a glycophorin A - glycophorin B fusion gene product that is detected using serology as the blood group antigen Dantu NE+. This DUP4 variant confers a clinically-important protective effect, with carriers ~40% less likely to develop cerebral malaria (5). Given that glycophorin A and glycophorin B are erythrocyte receptors for the *Plasmodium falciparum* cell surface receptors MSP-1, EBA-175 and EBL-1, respectively; it is likely that this protective effect is mediated by an altered ability of the parasite to invade the host erythrocyte (25–28).

A model for the DUP4 variant, based on analysis of mis-mapping positions of short-read sequences, was put forward that involved a duplication of the *GYPE* gene, deletion of the *GYPB* gene, and generation of two copies of a *GYPB-A* fusion gene (5). It is not clear how the GYPB-A fusion gene confers protection against malaria, but it has been suggested that the gene products could affect interactions with *Plasmodium falciparum* receptors and host band 3 protein at the erythrocyte surface. A complete characterization of the DUP4 allele is therefore important to understand the mechanistic basis for this protective effect, and to facilitate design of treatments mimicking the mechanism of protection.

## Results

### The physical structure of the DUP4 structural variant

We initially decided to confirm this structural model by physical mapping of the DUP4 variant using fiber-FISH. We grew lymphoblastoid cells from the known DUP4 heterozygote from the 1000 Genomes project sample collection, HG02554, derived from a man with African ancestry from Barbados, and selected clones to act as fiber-FISH probes from the WIBR-2 human fosmid library spanning the region, based on fosmid end sequences previously mapped to the GRChr37 reference genome.

Fiber-FISH showed that the reference haplotype generates signals consistent with the genome reference sequence (Figure 1a). Because of the high sequence identity between the tandemly-repeated glycophorin regions, there is extensive cross-hybridization of probes that map to the *GYPB* repeat with the *GYPE* and *GYPA* repeats. The *GYPE* repeat can be distinguished by hybridization of a small *GYPE-*repeat-specific PCR product, and the *GYPA* repeat can be identified by a gap in the green fosmid probe signal, caused by 16kb of unique sequence in the *GYPA* repeat. Also, the overlap of the distal end of the blue fosmid probe with the proximal end of the *GYPE* repeat allows a small amount of blue signal at the distal end of the *GYPB* and *GYPE* repeats, confirming repeat length and orientation. We identified DNA fibers showing an arrangement completely consistent with the DUP4 model proposed previously (Figure 1b).

**Figure 1.**
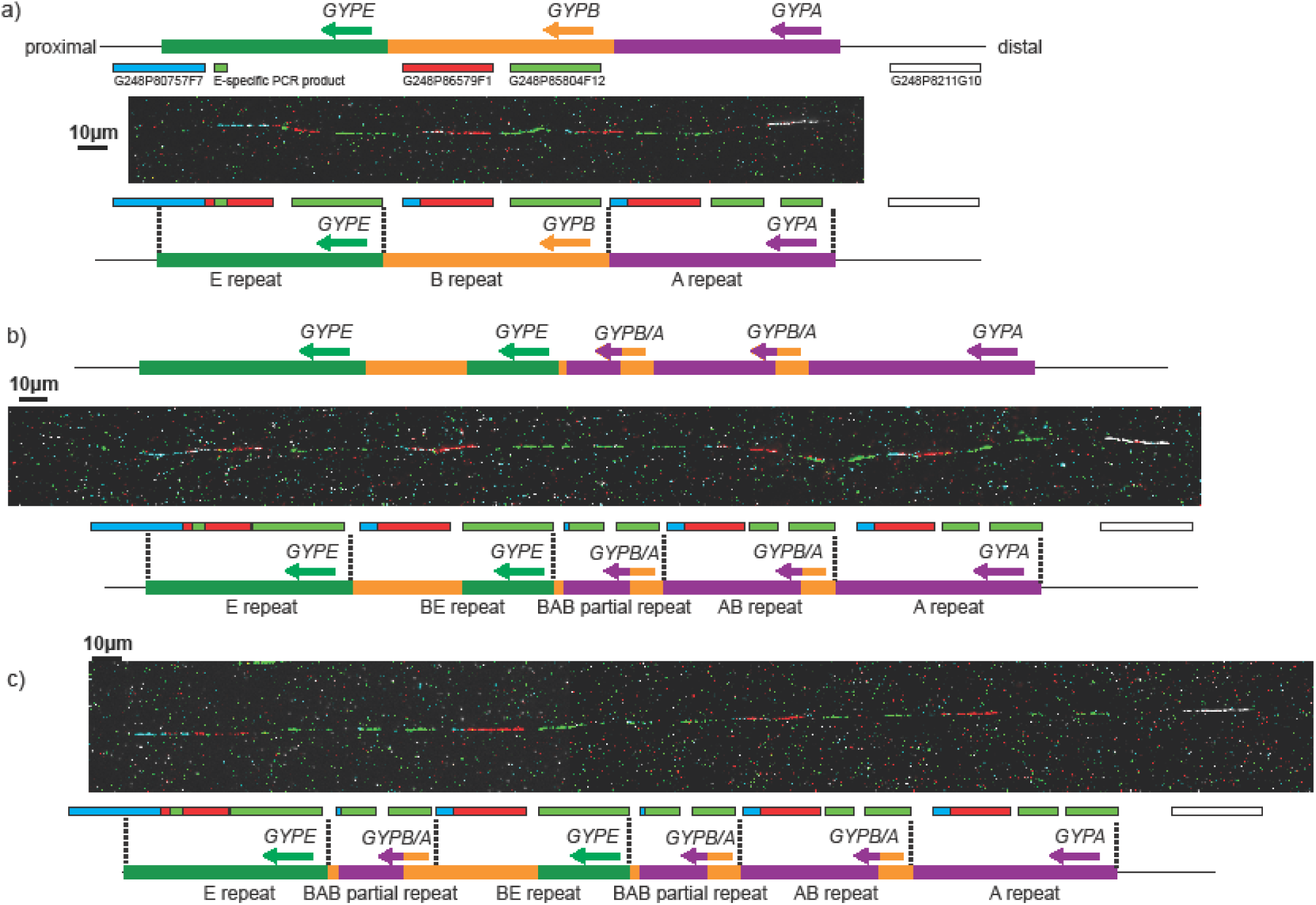
Fiber FISH analysis of the DUP4 heterozygote sample HG02554. a) An example DNA fiber from the reference haplotype. The position and label color of the fosmid probes is indicated above the fiber on a representation of the human reference genome, and the interpretation of the FISH signals shown below the fiber. b) An example DNA fiber from the DUP4a haplotype. The Leffler model of the DUP4 haplotype is indicated above the fiber. The interpretation of the FISH signals shown below the fiber. c) An example DNA fiber from the DUP4b haplotype. The interpretation of the FISH signals is shown below the fiber.

### Identification of a somatic DUP4 variant

However, we also visualized fibers with an extra partial repeat unit, which we called DUP4b (Figure 1c). This novel variant carries an extra copy of the partial A-B repeat, which harbors the *GYPB/GYPA* fusion gene. We selected a fosmid probe that spanned the 16kb insertion specific to the *GYPA* repeat, and showed that the extra copy was at least partly derived from the A repeat, consistent with the extra copy being an extra copy of the partial A-B repeat (Figure S1).

To rule out large-scale karyotype changes being responsible for our observations of the additional novel variant (DUP4b), we analyzed metaphase spreads of HG02554 lymphoblastoid cell line using metaphase-FISH, interphase FISH and multiplex-FISH karyotyping (Figure S2). DUP4 and reference chromosomes could be distinguished by interphase-FISH on the basis of hybridization intensity of a fosmid probe mapping to *GYPB* (Figure S2b). No evidence of large scale inter- and intrachromosomal rearrangements or aneuploidy was found in any of our experiments.

We hypothesized that DUP4b is a somatic variant that occurred through rearrangement of the original DUP4 variant (which we call DUP4a), but not the reference variant. If this is true, we would expect to observe an equal number of reference and DUP4 fibers from each of the parental chromosomes confirming the heterozygous DUP4 genotype of the source cells, but the DUP4 fibers to be subdivided into DUP4a and DUP4b variants. Of 24 fibers examined from HG02554, 12 were reference and 12 were DUP4, and, of the 12 DUP4 variants, 7 were DUP4a and 5 were DUP4b, strongly supporting the model where DUP4b is a somatic rearrangement of DUP4a and the presence of two sub-clones (populations) of cells, one with reference and DUP4a haplotypes, the other with reference and DUP4b haplotypes. We also analyzed the HG02554 cell line from the Oxford laboratory used in their study (5), and confirmed the existence of DUP4b by fiber-FISH. The high frequency of DUP4b variant chromosomes within the cell lines together with the observation of DUP4b in two cell line cultures suggests that DUP4b is a somatic variant of DUP4a that has arisen prior to the passage received by the Oxford laboratory or the Wellcome Trust Sanger Institute, either in the donor individual, or early in the cell-culturing process, perhaps increasing in frequency due to the associated transformation cell bottleneck (29).

To further characterize the somatic variation observed in HG02554, we Illumina sequenced at high depth (50x) HG02554 DNA purchased directly from Coriell Cell Repositories and extracted from their HG02554 lymphoblastoid cell line rather than extracted from our cell lines, together with peripheral-blood derived genomic DNA from two Tanzanian DUP4 homozygotes and three Tanzanian DUP4 heterozygotes. Analysis of sequence read depth across the glycophorin repeat region showed the same pattern to that observed previously (5), leading to a model that is confirmed by our fiber-FISH data (Figure 2a). DUP4 homozygotes show the expected increase to 4 copies and 6 copies in duplicated and triplicated regions, respectively.

**Figure 2.**
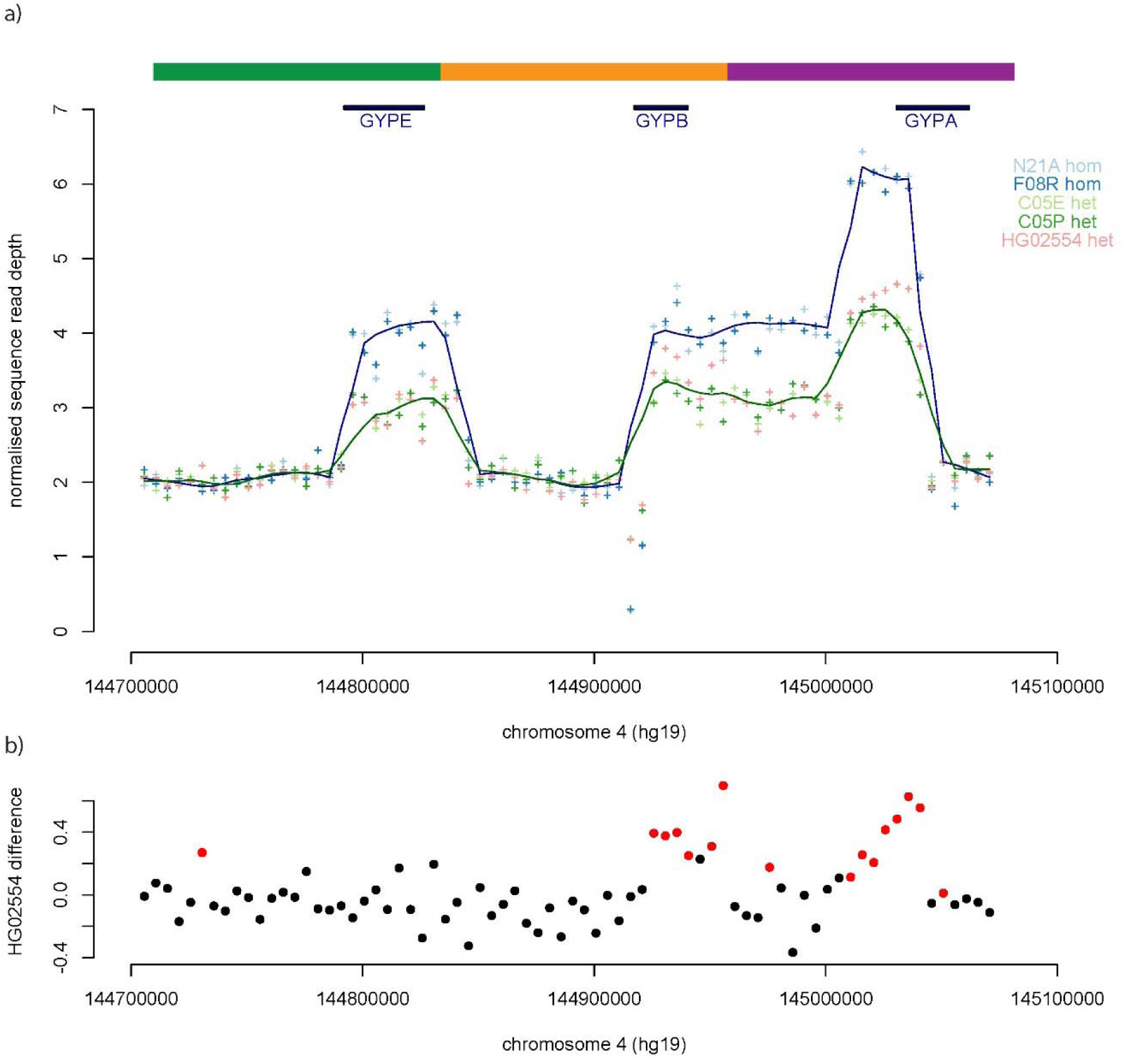
Sequence read depth analysis of DUP4 homozygotes and heterozygotes. a) Normalised sequence read depth of 5 kb windows spanning the reference sequence glycophorin region for five samples. The lines show the Loess regression line (f=0.1) for homozygotes (blue) and heterozygotes (green). Gene positions and repeats, with respect to the reference sequence, are shown above the plot. b) The difference in HG02554 sequence read depth compared to the average sequence read depth of the two other heterozygotes C05E and C05P is shown in 5kb windows across the glycophorin region. Points highlighted in red are significantly different (p<0.01).

We then compared the sequence read depth of HG02554 to the other two DUP4 heterozygotes to search for evidence of an increased copy number of the BAB partial repeat carrying the *GYPA/GYPB* fusion gene (Figure 1) suggested by our fiber-FISH data, which would reflect somatic mosaicism. HG02554 indeed shows a significant increase in DNA dosage in regions matching the BAB repeat, of around about 0.5, reflecting an extra copy of the region in ~50% of cells (Figure 2b).

### Development of a simple robust DUP4 genotyping assay

Having characterized the structure of DUP4 variants, we designed a simple robust junction fragment PCR assay that would allow detection of the DUP4 variant (both DUP4a and DUP4b) in nanograms of genomic DNA, at a large scale. This involved designing allele-specific and paralogue-specific PCR primers across a known breakpoint, a process made more challenging by the high sequence identity between paralogs. DUP4-specific primers had a modified locked nucleic acid base incorporated in the terminal 3’ nucleotide to enhance specificity for the correct paralog (30). We initially targeted the GYPA-GYPB breakpoint that created the fusion gene, but found that a similar breakpoint was present in a frequent gene conversion allele. However, we designed primers by targeting the breakpoint between the GYPE repeat and the GYPB repeat which was predicted to be unique to DUP4.

The DUP4 genotype was determined using a duplex PCR approach, with one pair of primers specific for the DUP4 variant, and a second pair amplifying across the SNP rs186873296, outside the structurally variable region, acting as a control for PCR amplification. The assay was validated against control samples showing different structural variants (5), and samples showing no structural variation, to ensure DUP4 specificity (Table S1, Figure S3).

### Association of DUP4 with malaria-related phenotypes

The DUP4 genotyping assay allowed us to investigate the association of DUP4 with three quantitative traits related to malaria: hemoglobin levels in peripheral blood, parasite load and mean number of clinical episodes of malaria, with hemoglobin levels showing the highest heritability of the three phenotypes in this cohort (Table 1). The DUP4 structural variant has previously been associated with both severe cerebral malaria and severe malarial anemia, and both are diagnoses related to our quantitative traits. For example, although the causes of hemoglobin level variation between individuals from a malaria-endemic region will be multifactorial, they will be strongly affected by malaria infection status of the individual, with infected individuals showing lower levels of hemoglobin (31). At the extreme low end of the distribution of hemoglobin levels is anemia, a sign of malaria which is one important feature in the pathology of the disease (32).

**Table 1.** Association of DUP4 allele with malarial phenotypes in the Nyamisiti cohort

| **Phenotype** | **Heritability of phenotype (95% CI)\*** | **Individuals** | **association p-value** |
| --- | --- | --- | --- |
| **Hemoglobin** | 0.302 (0.136-0.469) | 800 | 0.0054 |
| **Parasite load** | 0.104 (0.002-0.206) | 864 | 0.39 |
| **Clinical episodes** | 0.221 (0.131-0.311) | 939 | 0.72 |

\* calculated on this cohort using SOLAR (33, 34, 45)

We analyzed data from a longitudinal study of a population from the village of Nyamisati, in the Rufiji river delta, 150km south of Dar-es-Salaam, Tanzania, described previously (33–35). This region was holoendemic for malaria, predominantly *P. falciparum*, which causes 99.5% of all recorded clinical episodes of malaria. Parasite prevalence was recorded as 75% at the start of the study in 1993, falling to 48% in 1998, as measured by microscopy in the 2–9year old children. Previous work has suggested that the DUP4 variant is at a frequency of about 3% in the Wasambaa of north-eastern Tanzania (5), and our analysis found an allele frequency of 13.4% (95% confidence intervals 11.0%-16.1%) in 348 unrelated Tanzanians from Nyamisati in eastern coastal Tanzania, suggesting that this study would have power to detect a medium size of effect. A total of 962 individuals with pedigree information were genotyped. Using a family-based association method modelled in QTDT (36), we found a statistically significant association of the DUP4 variant with hemoglobin levels (p=0.0054, Table 1). We estimated the direction of effect by comparing the mean corrected hemoglobin levels of unrelated individuals with and without the DUP4 variant. Individuals with the DUP4 allele showed a higher hemoglobin level compared to those without a DUP4 variant, showing that DUP4 variant is associated with higher hemoglobin levels (Supplementary methods).

## Discussion

In summary, we directly demonstrate that the DUP4 variant has a complex structure involving a duplication of *GYPE*, deletion of *GYPB* and generation of two *GYPB/GYPA* fusion genes. The evolution of this particular rearrangement remains unclear. A model involving three intermediates has been suggested but none of these putative intermediates have yet been found (5). Given the relatively limited numbers of individuals analyzed for glycophorin structural variation so far, it is possible that these intermediate variants are rare or have been lost from the population. Indeed, given the extensive structural variation seen already at this locus, it seems likely that a high rate of genomic rearrangement generates complex variants which are mostly lost by genetic drift, with a few, such as DUP4, increasing in frequency due to positive selection. Further studies on the extensive variation in Africa are needed to fully characterize the variation at this locus.

We show that the DUP4 variant is associated with hemoglobin levels in a community setting indicating protection from malaria. Low levels of hemoglobin indicate anemia, which can reflect sub-clinical levels of malaria infection, and the village studied has a very high prevalence of *P. falciparum* infection, our study supports the importance of the DUP4 variant in malaria protection. However, the absence of an association with either the number of clinical episodes of malaria or the parasite load is perhaps more puzzling. This may reflect the lower heritability of these traits compared to hemoglobin levels, and therefore the increased effect of non-genetic variation (Table 1). How DUP4 protects against malaria is unknown and alternatively these results may point to a role in directly affecting erythrocyte invasion by the parasite, which is detectable in our cohort, rather than the more general phenotypes such as number of clinical malaria episodes or parasite load.

We also show that a novel somatic variant exists (DUP4b) with an extra *GYPB/GYPA* fusion gene, suggesting that this region may be prone to somatic rearrangements. We cannot rule out a somatic rearrangement in the transformation and culturing of the lymphoblastoid cell line, although it has been shown previously that such genetic changes introduced by EBV transformation are either rare (15, 37), or overlap with regions known to undergo extensive programmed somatic rearrangement, such as the immunoglobulin loci (38). It is possible, therefore, that the somatic variant originated in the donor patient given that the HG02554 B-lymphoblastoid cells from Oxford and the Wellcome Trust Sanger Institute were both from the same batch of cells (passage #4, according to Coriell Cell Repositories); recent evidence suggest that such structural variant mosaics are likely to occur at a significant frequency, at least at certain loci (39, 40). We demonstrate that this somatic variant is able to be detected from high coverage short read sequence data, which will allow further analysis of somatic variation at this locus without cell material. Our data raises the intriguing possibility of heightened somatic instability and somatic mosaicism at this locus in DUP4 carriers, which might confer added protection against malaria.

## Methods

### Study population, samples and phenotypic data

The study population is from the coastal village of Nyamisati in the Rufiji delta in Tanzania (35). Human genomic DNA was extracted from whole peripheral blood, as previously described, with informed consent and approval of the local ethics committees of the National Institute of Medical Research in Tanzania and the Regional Ethical Committee of Stockholm in Sweden.

Phenotypic data spanning 7 years, from 1993 to 1999, was collected using annual total population surveys and annual records of malarial episodes, as previously described (34). The total population survey provided information on asymptomatic parasite load (parasites per μl), and hemoglobin levels. A single hemoglobin value was generated for each individual from annual total population surveys carried out over 7 years, corrected for age and sex and parasite load prior to genetic analysis. A single parasite load (parasites per μl) was generated for each individual from asymptomatic parasitemia recorded during annual total population surveys carried out over 7 years. This single figure was corrected for age and sex prior to genetic analysis.

All clinical malarial episodes were recorded, and confirmed by microscopy. Multiple clinical episodes were recorded if the recurrence was greater than 4 weeks apart. A small proportion (1%) of individuals presented with a clinical episode at the time of the total population survey, and these samples were included in the analysis. A single clinical episodes value was generated for each individual from records of all malarial episodes occurring in the village during a period of 7 years. The phenotypic data derived for clinical episodes, parasite load and hemoglobin has previously been described in detail (33, 34).

### Fluorescence *in situ* hybridization using single-molecule DNA fibers (Fiber-FISH)

The probes used in this study included four fosmid clones selected from the UCSC Genome Browser GRCh37/hg19 assembly and a 3632-bp PCR product that is specific for the glycophorin E repeat (see below). Probes were made by whole-genome amplification with GenomePlex Whole Genome Amplification Kits (Sigma-Aldrich) as described previously (41). Briefly, the purified fosmid DNA and the PCR product were amplified and then labeled as follow: G248P86579F1 and glycophorin E repeat-specific PCR product were labeled with digoxigenin-11-dUTP, G248P8211G10 was labeled with biotin-16-dUTP, G248P85804F12 was labeled with DNP-11-dUTP and G248P80757F7 was labeled with Cy5-dUTP. All labeled dUTPs were purchased from Jena Bioscience.

The preparation of single-molecule DNA fibers by molecular combing and fiber-FISH was as previously published (42) (3), with the exception of post-hybridization washes, which consisted of three 5-min washes in 2× SSC at 42°C, instead of two 20-min washes in 50% formamide/50% 2× SSC at room temperature.

### Interphase-, metaphase-FISH and karyotyping by multiplex-FISH

Metaphase chromosomes were prepared from a human lymphoblastoid cell line (HG02554) purchased from Coriell Cell Repositories. Briefly, colcemid (Thermo Fisher Scientific) was added to a final concentration of 0.1 μg/ml for 1 h, followed by treatment with hypotonic buffer (0.4% KCl in 10 mM HEPES, pH7.4) for 10 min and then fixed using 3:1 (v/v) methanol:acetic acid.

For interphase- and metaphase-FISH, G248P8211G10 labeled with Texas Red-dUTP, G248P85804F12 labeled with Atto488-XX-dUTP (Jena Bioscience), and RP11-325A24 labeled with Atto425-dUTP (Jena Bioscience) were used as probe. Slides pre-treatments included a 10-min fixation in acetone (Sigma-Aldrich), followed by baking at 65°C for 1 hour. Metaphase spreads on slides were denatured by immersion in an alkaline denaturation solution (0.5 M NaOH, 1.0 M NaCl) for 10 min, followed by rinsing in 1M Tris-HCl (pH 7.4) solution for 3 min, 1× PBS for 3 min and dehydration through a 70%, 90% and 100% ethanol series. The probe mix was denatured at 65°C for 10 min before being applied onto the denatured slides. Hybridization was performed at 37°C overnight. The post-hybridisation washes included a 5-min stringent-wash in 1× SSC at 73°C, followed by a 5-min rinse in 2× SSC containing 0.05% Tween®20 (VWR) and a 2-min rinse in 1× PBS, both at room temperature. Finally, slides were mounted with SlowFade Gold^®^ mounting solution containing 4’6-diamidino-2-phenylindole (Thermo Fisher Scientific). Multiplex-FISH (M-FISH) with human 24-color painting probe (43).

Slides were examined using AxioImager D1 microscope equipped with appropriate narrow-band pass filters for DAPI, Aqua, FITC, Cy3 and Cy5 fluorescence. Digital images capture and processing were carried out using the SmartCapture software (Digital Scientific UK). Ten randomly selected metaphase cells were karyotyped based on the M-FISH and DAPI-banding patterns using the SmartType Karyotyper software (Digital Scientific UK).

### PCR for fiber-FISH probe generation

The 3632 bp glycophorin E repeat-specific PCR product for use as a fiber-FISH probe was generated by long PCR. Long PCRs were performed in a total volume of 25μl using a *Taq/Pfu* DNA polymerase blend (0.6U *Taq* DNA polymerase/ 0.08U *Pfu* DNA polymerase), a final concentration of 0.2μM primers Specific_glycophorinE_F and and Specific_glycophorinE_R (Table S2), in 45mM Tris-HCl(pH8.8), 11mM (NH_4_)_2_SO_4_, 4.5mM MgCl_2_, 6.7mM 2-mercaptoethanol, 4.4mM EDTA, 1mM of each dNTP (sodium salt), 113μg/ml bovine serum albumin. Cycling conditions were an initial denaturation of 94°C for 1 min, a first stage consisting of 20 cycles each of 94°C for 15s and 65°C for 10 min, and a second stage consisting of 12 cycles each of 94°C for 15 s and 65°C for 10 min (plus 15s/cycle); these were followed by a single incubation phase of 72°C for 10 min.

### Illumina sequencing of DUP4 samples

1 μg genomic DNA was randomly fragmented to a size of 350bp by shearing, DNA fragments were end polished, A-tailed, and ligated with the NEBNext adapter for Illumina sequencing, and further PCR enriched by P5 and indexed P7 oligos. The PCR products were purified (AMPure XP system) and theresulting libraries were analyzed for size distribution by an Agilent 2100 Bioanalyzer and quantified using real-time PCR.

Following sequencing on a Illumina platform, the resulting 150bp paired-end sequences were examined for sequencing quality, aligned using BWA to the human reference genome (hg19 plus decoy sequences), sorted using samtools (44) and duplicate reads marked using Picard, generating the final bam file. Sequencing and initial bioinformatics was done by Novogene Ltd.

Sequence read depth was calculated in using samtools to count mapped reads in non-overlapping 5kb windows across the glycophorin region. Read counts were normalised for coverage to a non-CNV region (chr4:145516270-145842585), then to the first 50kb of the glycophorin region which has diploid a copy number of 2.

### DUP4 junction fragment PCR genotyping

Primer sequences are shown in table S2. PCR was conducted in a final volume of 10μl in 1× Kapa A PCR buffer (a standard ammonium sulfate PCR buffer) with a final concentration of 1.5mM MgCl_2_, ~10ng genomic DNA, 0.2mM of each of dATP, dCTP, dGTP and dTTP, 1U *Taq* DNA Polymerase, 0.1μM each of rs186873296F and rs186873296R, and 0.5μM each of DUP4F2 and DUP4R2. Thermal cycling used an ABI Veriti Thermal cycler with an initial denaturation of 95°C for 2 minutes, followed by 35 cycles of 95°C 30 seconds, 58°C 30 seconds, 70°C 30 seconds, then followed by a final extension of 70°C for 5 minutes. 5ul of each the PCR products were analyzed using standard horizontal electrophoresis on an ethidium-bromide-stained 2% agarose gel.

Routine genotyping included a DUP4 positive control in every experiment (sample HG02554). We distinguished homozygotes by quantification of DUP4 and control band intensity on agarose gels using ImageJ, and calculating the ratio of DUP4:control band intensity for each individual. At low allele frequencies, homozygotes are expected to be rare. After log2 transformation, a cluster of four outliers of high ratio (>2SD, log2ratio>1.43) were clearly separated from the 278 other DUP4 positive samples, and these four were classified as homozygotes. The remaining 278 DUP4 positive samples were classified as heterozygotes.

### Family-based association analysis

Associations between the three clinical phenotypes and DUP4 genotype were tested using QTDT version 2.6.1 (36) on the full dataset of 167 pedigrees, using an orthogonal model. The heritability for all the clinical phenotypes was initially estimated using a model for polygenic variance. A test for total evidence of association was performed which included all individuals within the samples, retaining as much information as possible. This total test of association included environmental, polygenic heritability and additive major locus variance components within the model. To control for population stratification within family association was tested in an orthogonal model including environmental, polygenic heritability and additive major locus variance components. Direction of effect was estimated by comparing the hemoglobin level values, expressed as residuals from the regression model used to correct for age and sex, between the 262 unrelated individuals with hemoglobin level values from the Nyamisati cohort carrying (n=70, mean=1.67 g/L, standard deviation=16.0 g/L) and not carrying (n=192, mean=−0.12 g/L, standard deviation=14.1g/L) the DUP4 variant.

## Acknowledgements

This work was funded by a SACB PhD studentship to WA, and grant WT098051 (FY & SL). We thank Ellen Leffler and Gavin Band for helpful comments on a previous version of this manuscript, Kirk Rockett for providing the HG02554 cells used by the Oxford laboratory, and Chris Tyler-Smith for support.

## Author contributions

The study was designed and conceived by EJH. Experimental work was carried out by WA, SL, PB, FY and EJH. Data analysis was carried out by DC, WA and EJH. Malaria phenotype data and samples were provided by AF, DC, IR and M-AS. The paper was written by EJH with contributions from all authors. All authors approved the final manuscript.

**Figure S1.**
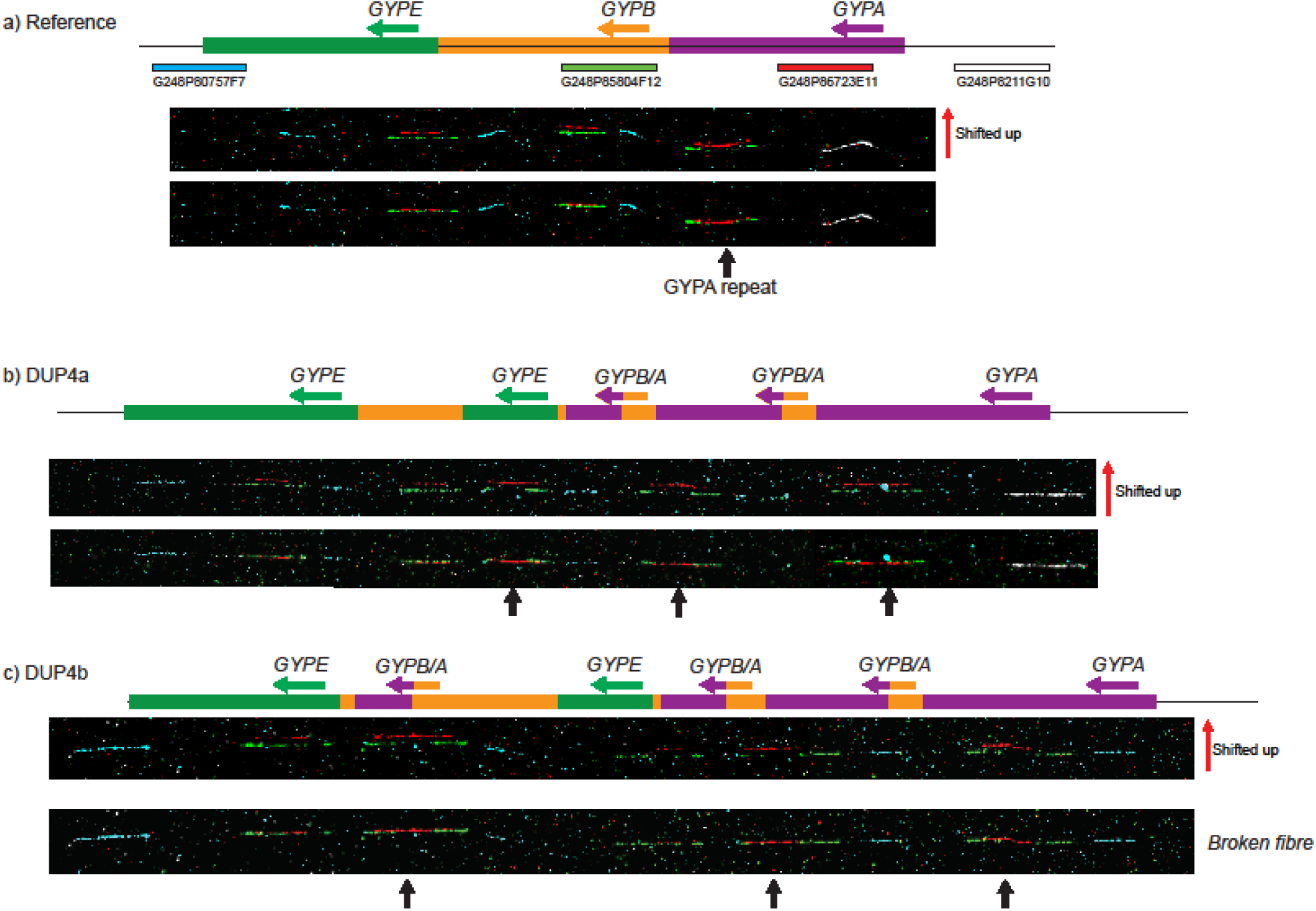
Fiber FISH analysis of the DUP4 heterozygote sample HG02554 using a GYPA-repeat fosmid probe. a) An example DNA fiber from the reference haplotype. The names, relative position and colors of the fosmid probes are indicated above the fiber. The upper image is generated from the lower image, shifting up the red signal to show the gap in the green fosmid signal specific to the GYPA repeat. b) An example DNA fiber from the DUP4a haplotype. The Leffler model of the DUP4 haplotype is indicated above the fiber, and arrows indicate GYPA repeat units or GYPA partial repeat units. Note that the complete DUP4 pattern consists of two images of the same fiber stitched together. c) An example DNA fiber from the DUP4b haplotype. Our model of the DUP4b somatic variant haplotype is indicated above the fiber, and arrows indicate GYPA repeat units or GYPA partial repeat units. Note that for this experiment although several images were captured, none were from a full-length fiber with both distal (white) and proximal (blue) fosmid signals. In this fiber, the white and GYPA signals were missing from the distal end of the fiber.

**Figure S2.**
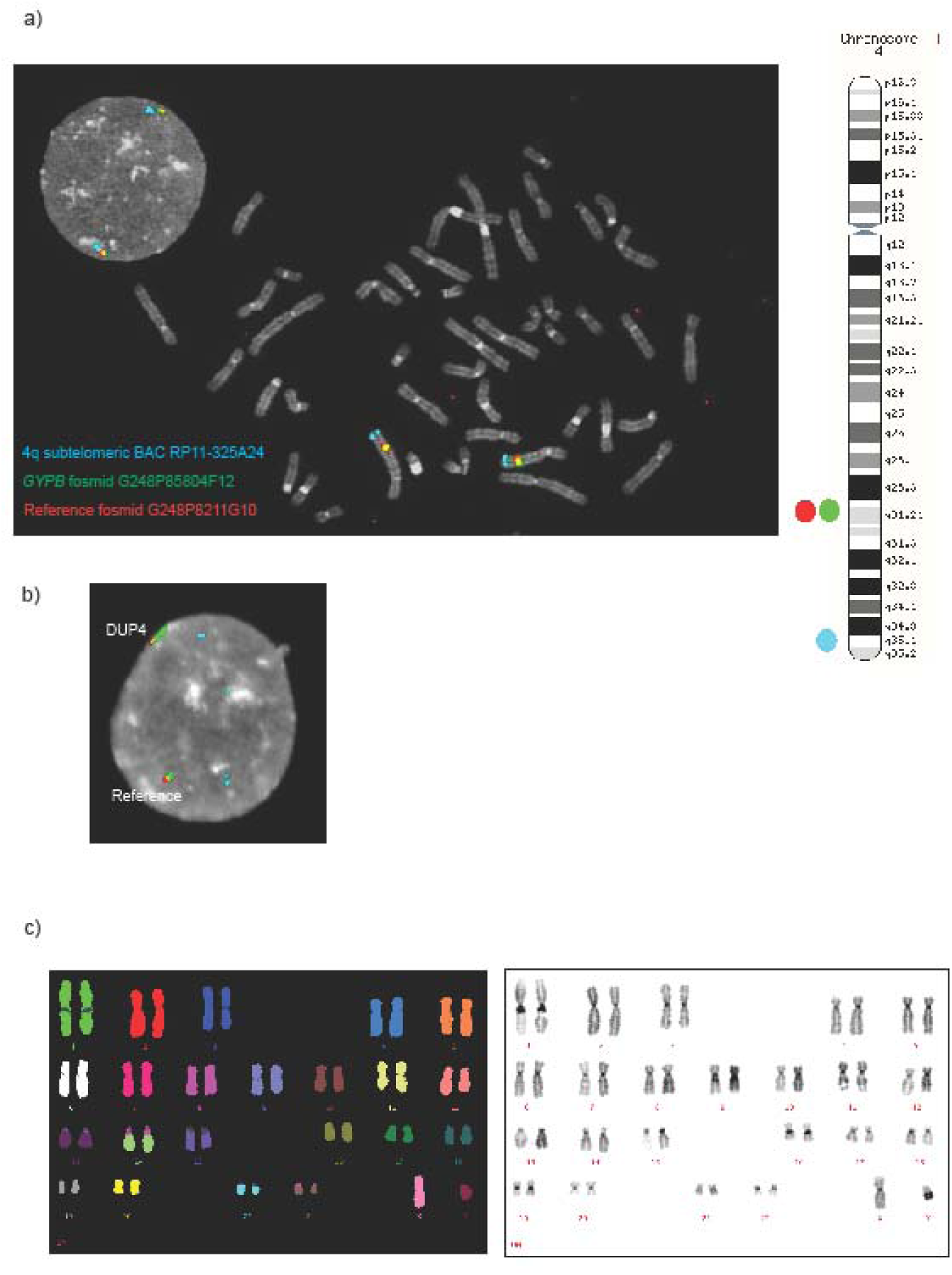
FISH analysis of the DUP4 heterozygote sample HG02554. a) Metaphase-FISH and interphase-FISH analysis of the glycophorin region. Probe locations shown in the ideogram on the right, and shown in more detail in figure 1. b) An interphase-FISH magnified image, showing that the DUP4 variant can be distinguished by a larger signal from the green probe mapping to the GYPB repeat. c) Multiplex FISH (left) and DAPI-stained banding (right) of HG02554 cell line, showing no major rearrangements or aneuploidies.

**Figure S3.**
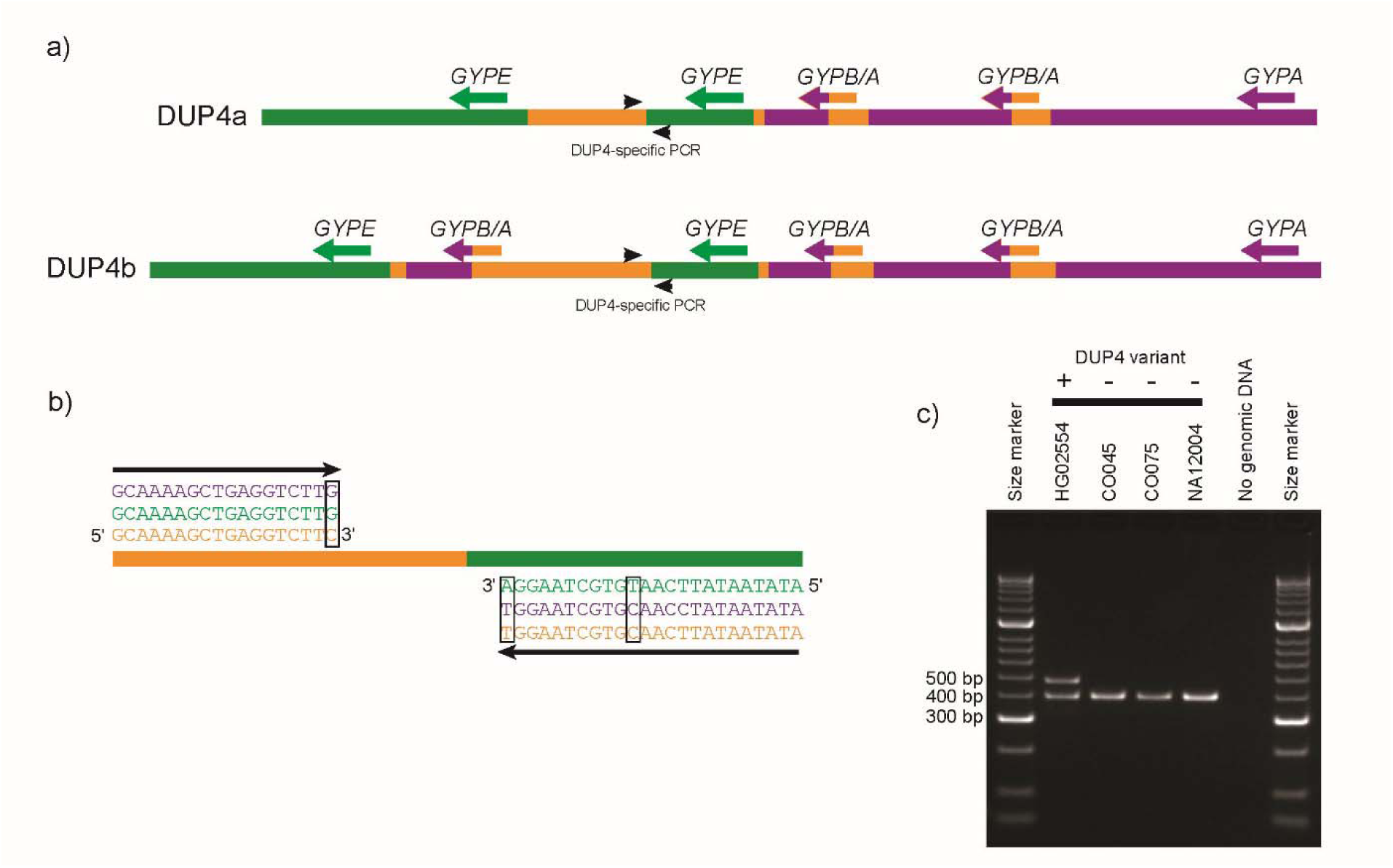
Genotyping using DUP4 junction fragment PCR. a) PCR primer design strategy for a DUP4-specific PCR b) PCR primers incorporating mismatches to ensure paralogue specificity across the B-repeat/E-repeat breakpoint. c) An image of an ethidium bromide-stained agarose gel showing PCR products generated by the DUP4 junction fragment PCR (~500bp) and a co-amplified control PCR product (~400bp). A known DUP4 heterozygous positive sample (HG02554) and three DUP4 negative samples are shown.

## SUPPLEMENTARY MATERIAL

**Table S1.** Control samples for DUP4 genotyping assay

| Sample (1000 genomes) | Population | Structural variant carried (reference 8) | DUP4 assay result |
| --- | --- | --- | --- |
| HG02554 | ACB | DUP4 | Positive |
| NA18646 | CHB | DUP14 | Negative |
| NA19084 | JPT | DUP28 | Negative |
| HG02679 | GWD | DUP7 | Negative |
| NA18625 | CHB | DUP2 | Negative |
| NA19360 | LWK | DUP3 | Negative |
| NA19474 | LWK | DUP3 | Negative |
| HG02716 | GWD | DEL7 | Negative |
| NA19084 | JPT | DUP28 | Negative |
| NA19085 | JPT | None | Negative |
| NA19190 | YRI | DEL2 | Negative |
| NA19777 | MEX | None | Negative |
| NA19818 | ASW | DEL2 | Negative |
| HG02585 | GWD | DUP5 | Negative |
| NA19120 | YRI | DEL1 | Negative |
| HG04039 | STU | DEL6 | Negative |
| HG03837 | STU | DUP24 | Negative |
| NA12004 | CEU | None | Negative |
| NA11839 | CEU | None | Negative |
| NA07034 | CEU | None | Negative |
| NA07055 | CEU | None | Negative |
| NA12814 | CEU | None | Negative |
| NA12005 | CEU | None | Negative |

**Table S2.**
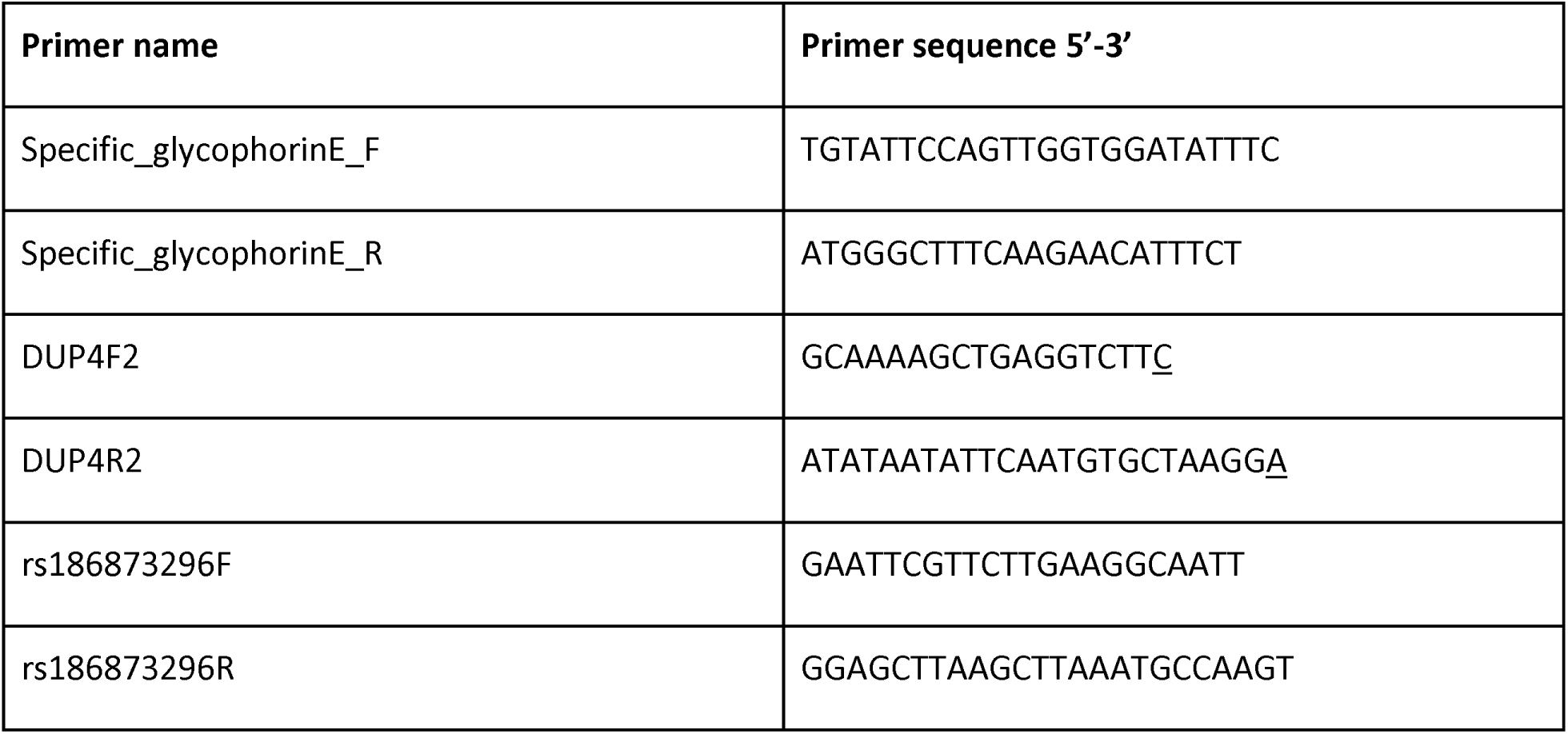
Primer sequences used in this study. Underlined nucleotides indicate that a linked nucleic acid (LNA) nucleotide was used at that position to increase paralog-specificity.

